# Iron-deficiency in the tumor microenvironment reprograms tumor-immune interactions in a sex biased manner

**DOI:** 10.64898/2026.05.29.728615

**Authors:** Aurosman Sahu, Yue Hao, Angela Baker, Kondaiah Palsa, Tim Helmuth, Ganesh Shenoy, Elizabeth B. Neely, Becky Slagle-Webb, Michael E. Berens, James Connor

## Abstract

**Background:** Glioblastoma (GBM) is a sex-biased disease characterized by higher incidence and poorer survival in males. These sex differences are primarily driven by metabolic and immune signatures, with iron metabolism playing a major role. While iron is essential for tumor cell proliferation, it is also critical for T cell recruitment and function within the tumor microenvironment (TME). Clinical data indicates that iron deficiency impacts GBM survival in a sex-biased manner; however, the underlying mechanisms remain unexplored.

**Methods:** We employed a FTH1 heterozygous knockdown mouse model to induce tumor iron deficiency in GBM-bearing mice. TME dynamics were interrogated using flow cytometry and spatial transcriptomics (10X Xenium) to analyze immune infiltration, localization, and ligand-receptor signaling between GBM and immune cells.

**Results:** FTH1 knockdown resulted in iron deficiency in the GBM TME. Iron deficiency altered the TME dynamics in a sex biased manner. FTH1 knockdown in females caused an anti-inflammatory, cytokine deficient TME which failed to recruit CD4 and CD8 T cells. In males, FTH1 knockdown caused a proinflammatory environment by activating the innate immune response. Additionally, FTH1 knockdown increased the density of TAMs in the immediate surrounding of GBM cells in a sex biased manner.

**Conclusion:** These results demonstrate that tumor iron plays a sex-biased role in immune infiltration and anti-tumor immunity.

## Introduction

Glioblastoma (GBM) is one of the most aggressive brain malignancies, with a 5-year survival rate of 6 percent[1]. It is also extremely difficult to treat because of its high cellular heterogeneity, rapid invasiveness, and ability to evade treatments[2]. GBM cells are astrocytic in origin and characterized by a unique immunosuppressive microenvironment[3]. The tumor progression and response to therapy in GBM is driven by tumor intrinsic factors such as tumor metabolic signature and genetic mutations as well as extrinsic factors like infiltrating immune cells, brain resident non-tumor cells and endothelial cells[4]. Recent evidence suggests that GBM is a sexually dimorphic disease, where males tend to develop GBM more frequently and have worse survival[5]. These sex differences are driven by the combined effects of sex hormones and sex chromosomes, which can influence the epigenetic landscape, metabolic signatures, antitumor immunity and response to therapy[6]. Of all the metabolic processes, iron metabolism shows the highest degree of sex differences and has been linked with changes in GBM survival, TME dynamics and immune function[7-10].

Iron plays a vital role in human physiology due to its active involvement in oxygen transport. Iron is essential for several cellular processes, such as mitochondrial ATP production, DNA synthesis, and cell proliferation[11]. GBM cells have a high iron requirement due to their rapid proliferation rates, and they upregulate the transferrin receptor (TFRC), a protein involved in iron uptake [12]. TFRC mainly binds transferrin, a protein involved in iron transport that carries two iron atoms per protein molecule[13]. However, recent studies have shown that ferritin, typically considered an iron storage protein, also binds to TFRC [14, 15]. Studies from our laboratory and others have shown that H-Ferritin (FTH1), a subunit of ferritin, can act as an iron delivery protein and deliver iron to brain as well as chemotoxins [16, 17]. Hence, FTH1 mediated iron delivery can act as a major mode of iron uptake by GBM cells. Studies in GBM patients have shown that GBM tissues bind FTH1 in a sex biased manner (female GBM has greater binding than male), further consolidating the role of FTH1 mediated iron delivery in GBM progression and driving sex biased differences[18].

Iron plays an essential role in tumor growth, immune infiltration, TAM polarization and the overall dynamics in the TME[10, 19]. In GBM, iron metabolism contributes to sex biased differences in survival, tumor growth and immune infiltration[7, 8]. Because tumors rely heavily on iron to sustain rapid growth, altering iron availability within the tumor microenvironment represents a promising therapeutic strategy. In particular, iron chelators have been explored as potential anticancer agents, as they limit tumor proliferation by sequestering iron required for cellular proliferation.[20]. Iron deficiency in tumors can act as a double-edge sword. Iron deficiency in some cancers can promote tumor progression through activation of hypoxic responses, metabolic reprogramming, and suppression of the immune response, as well as in other cancers inhibit tumor growth by limiting tumor cell proliferation[21]. In GBM, iron deficiency has been shown to decrease patient survival in a sex biased manner[9]. However, the underlying mechanisms that drive these outcomes remain unexplored. Here, we used a genetic model to induce iron deficiency in GBM bearing mice to interrogate how tumor iron levels affect the TME dynamics. Previous research using the current model has shown that FTH1 knockdown decreases tumor bearing mice survival in a female biased manner[22]. Due to the closely intertwined nature of iron metabolism with immune response, we used flow cytometry and spatial transcriptomics to interrogate how iron deficiency in the tumor affects immune infiltration, localization and GBM-immune cell interactions in the TME.

## Methods

### Cell culture

GL261 (RRID: CVCL_Y003) cells were obtained from the Developmental Therapeutic Program, NCI. GL261 cells were cultured in DMEM with GlutaMAX (ThermoFisher Scientific, Catalog #: 61870036) with 10% fetal bovine serum (GeminiBio, Catalog number: 100–106), and 1% Penicillin-Streptomycin (ThermoFisher Scientific, Catalog #: 15140-122). Cells were maintained in a humidified tissue culture incubator with 5% CO2 at 37 °C. For experiments, cells up to passage number 8 post-thawing were included to maintain reproducibility. All cell lines tested negative for mycoplasma contamination.

### FTH1 Knockdown mice model and intracranial injection of GL261 cells

Since FTH1 homozygous knockout is embryonically lethal in mice, a heterozygous FTH1 knockdown mouse model was developed where the knockdown animals showed significant reduction in serum FTH1[23]. Mice of age 6-8 weeks were used for intracranial injections. Intracranial injections were performed using an adapted version of previously reported protocols[8, 22]. Briefly, six-week-old mice were anesthetized using inhaled isoflurane, and a Hamilton syringe attached to a stereotaxic apparatus was used to inject cells into the brain (-1.5mm lateral, 0.5mm rostral from the bregma) at a depth of approximately 3.5 mm. Each syringe was prepared with 2*10^4 GL261 cells suspended in 4 μL of null DMEM media. Animals were monitored over time for the presentation of neurological and behavioral symptoms. After 2 weeks post-injection, the tumor formation was confirmed by MRI imaging. All animal experiments were performed in compliance with institutional guidelines and were approved by the Institutional Animal Care and Use Committee (IUCAC) of the Penn State College of Medicine.

### Immune phenotyping of the tumors

At 3 weeks post-injection, immune cell profiling was performed on the mice bearing tumors. The mice were anesthetized using an intraperitoneal injection of Ketamine (100□mg/kg) and Xylazine (10□mg/kg). Once fully anesthetized as assessed by an absent toe-pinch reflex, mice were euthanized following a transcardial perfusion using a lactated ringer solution. The tumor bearing hemisphere was isolated from the brain and processed further to interrogate the immune cell populations. Single-cell suspensions were prepared from the tumor-bearing hemisphere by enzymatic digestion using collagenase IV (Sigma) and DNase I (Sigma), followed by filtering through a 70-µm cell strainer. The lymphocyte fraction was isolated using a discontinuous Percoll density gradient (70% Percoll + 35% Percoll)[24]. The cells were then treated with live/dead staining (Sigma) and aliquots incubated in fluorophore conjugated antibodies (CD45, CD11b, CD3, CD4 and CD8). Stained cells were imaged in a flow cytometer (BD FACSymphony), and the acquired data were analyzed with FlowJo software (v11, BD Biosciences).

### Spatial transcriptomics sample preparation

Following perfusion euthanasia, tumor-bearing hemispheres were prepared following the Xenium In Situ - Fresh Frozen Tissue Preparation Handbook. Briefly, tumor bearing brains were fixed with paraformaldehyde and cryoprotected in 30% sucrose for fresh frozen sectioning. One animal was used from each group for this analysis. The sections were mounted on 10X Xenium slide. Xenium slides with samples were processed using a predesigned 5K mouse panel of probes on the Xenium Analyzer (10X Genomic). Tissue staining was performed using a combination of nucleus (DAPI), cellular content, and cell boundary markers with the Xenium Multi-Tissue Stain Mix (10X Genomics). This staining delineates cell boundaries and allows the segmentation of cells throughout the tissue.

### Xenium downstream data analysis

Xenium data output was loaded on Xenium Explorer for general data visualization and analysis. For unsupervised clustering of the dataset, Xenium output data was processed using the Scanpy (v1.10.4) and Seurat packages. Harmony (harmonypy, v0.0.10) was used for batch correction and the clustering of the cells belonging to specific genotypes was done by linear discriminant analysis (LDA) with a used split point.

### Chemokine expression in the tumor

Tumor regions of interest (ROIs) were manually delineated on the spatially registered transcriptomic image using Napari (v0.6.6). To assess bulk chemokine gene expression within the tumor compartment, raw transcript counts were normalized to 10,000 counts per cell using scanpy and log-transformed. For each chemokine gene, bulk expression across all cells within each tumor group was aggregated, and expression levels were standardized by computing Z-scores across tumor groups, defined as the mean-centered, standard-deviation-scaled expression value for each gene. Z-score normalization was applied across the tumor group axis to enable cross-group comparison of relative chemokine enrichment independent of absolute transcript abundance.

### COMMOT and cellchat

The cell-cell communication probabilities were determined using the published protocol [25]. The GBM cells (cluster 0 and cluster 1) were annotated as sender cells and the T cells were annotated as receiver cells. A distance threshold of 100 microns was chosen for the analysis, and the ligand-receptor (LR) pairs were obtained from the cellchat mouse database. CellChat analysis was in R using the mice LR database[26].

### Proximity analysis between GBM and TAMs

To quantify the spatial relationship between immune cell populations and tumor cells, cell centroids (x,y coordinates) were extracted from the spatial transcriptomic metadata. Spatial infiltration was defined as the Euclidean distance from each individual immune cell to its nearest malignant neighbor. This was calculated using a k-nearest neighbor algorithm (via the FNN package in R). The resulting distribution of distances was visualized using kernel density estimation (KDE) plots, facilitating a comparison of “infiltration depth” between groups.

### Tumor iron measurements

The tumor bearing tissue was prepared by nitric acid digestion for ICP-AES (Inductively Coupled Plasma Atomic Emission Spectroscopy) analysis, and the amount of ^56^Fe was quantified using previously reported protocol [27]. Concentrations of ^56^Fe were normalized to tumor tissue weight to get normalized iron levels in the tumor tissues.

## Statistics

For flow cytometry and ICP-AES data analysis, one way ANOVA was used. The data were expressed as mean□±□SD and the statistical analysis was done using GraphPad Prism 9 (RRID: SCR_002798). One-way ANOVA followed by Tukey’s post hoc analysis test was used to detect statistical significance (p□<□0.05) between the multiple groups. The significance annotations are * = P□≤□0.05, ** = P□≤□0.01, *** = P□≤□0.001 for all the figures.

## Results

### Global knockdown of FTH1 in the host reduces the tumor iron levels in the GBM tissue

To interrogate the role of tumor extrinsic FTH1 levels in GBM, we used a FTH1 heterozygous knockout mouse model. Three weeks post intracranial injection of GL261 cells, the tumors were harvested (Fig 1A). Measured tissue iron levels in the tumors represented whether global extrinsic FTH1 knockdown changed the tumor iron levels. ICP-AES analysis of the tumor bearing tissue indicated that the tumors in FTH1 KD mice had lower iron levels than WT (Fig 1B). This was observed in both male and female mice, although in females the decrease was more prominent. Next, we also probed the expression of several genes which are designated markers for tissue iron status. Aco1 and Ireb2, which code for Iron regulatory proteins (IRP1/2), were upregulated in FTH1 KD. IRP1/2 upregulation is a classic marker for iron deficiency. Similarly, the tumor tissue expressions of Hjv (Hemojuvelin), Hfe (Homeostatic Iron Regulator) and Slc40a1 (FPN) were all decreased in both male and female FTH1 KD mice, indicating an iron deficiency cellular state in the tumors (Fig 1C). Overall, these data confirm that global heterozygous knockout of FTH1 in the host decreases GBM tumor iron levels.

**Fig 1:**
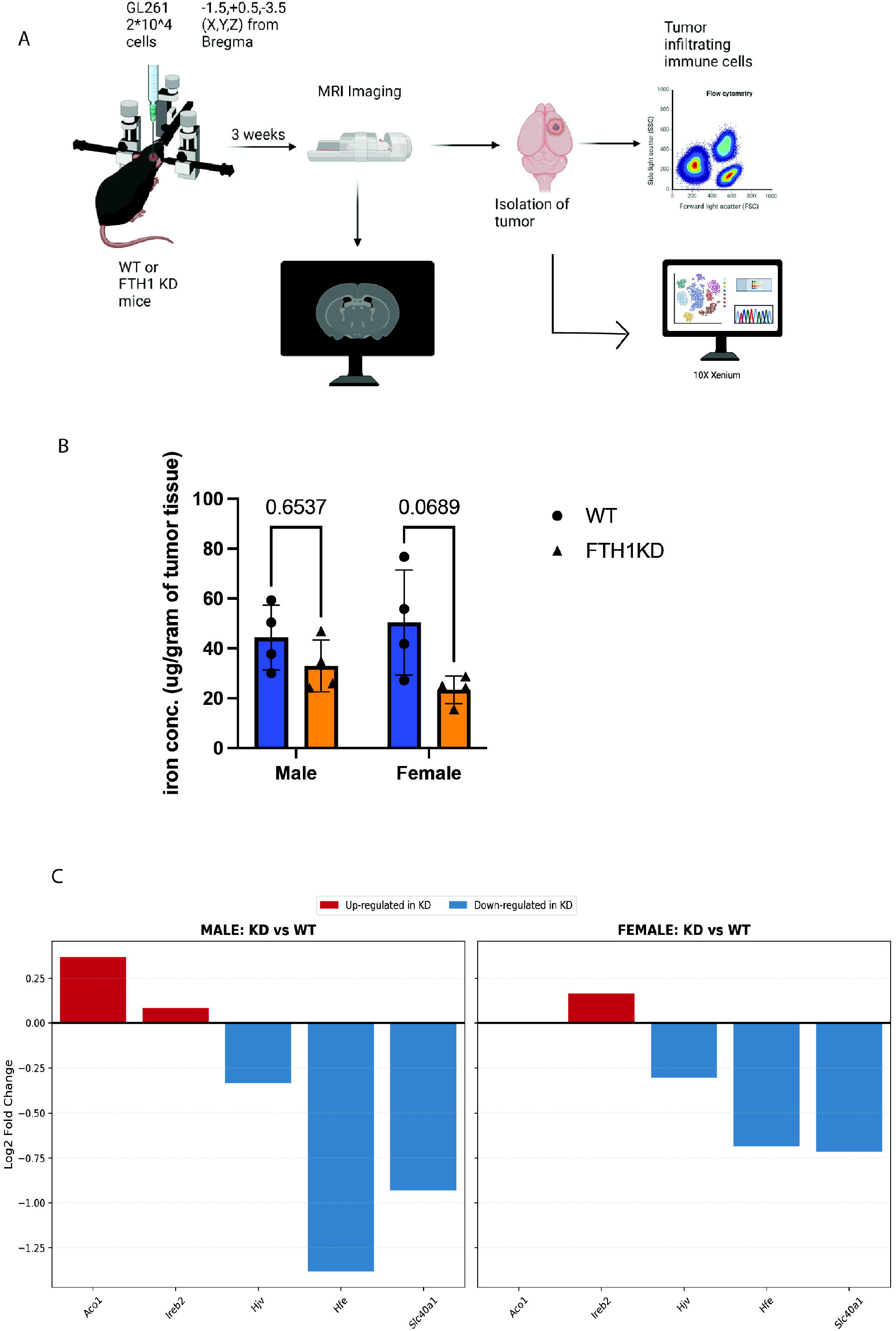
Systemic FTH1 depletion induces iron deficiency in GBM tissue. (A) Schematic representation of the mouse GBM model and downstream analysis. (B) ICP-AES showing extrinsic FTH1 knockdown causes a reduction of tumor iron levels (N = 4, One way ANOVA). (C) Relative mRNA expression of key iron marker genes indicating iron deficiency like signature in the tumor. (N = 1, Spatial Transcriptomics)

### Spatial transcriptomics analysis of tumor bearing brains

To further interrogate how extrinsic FTH1 knockdown affects GBM TME dynamics, we utilized the 10X Xenium platform with a 5K custom gene panel. We analyzed four tumor-bearing brains on a single slide, representing one of each genotype (WT vs. FTH1 KD) for each sex. H&E imaging confirmed the presence of tumors in the four brain tissues used for analysis (Fig 2A). Next, we assessed cell density across the tissues and found that the designated tumor regions in the H&E images corresponded precisely with areas of higher cell density (Fig 2B). We applied Linear Discriminant Analysis (LDA) to cluster cells by genotype, yielding four distinct clusters representing the four tissues (Fig 2C). Quantification of total cells per cluster revealed that the MKD brain hemisphere had the highest cell count, consistent with its larger tumor volume. The other three tumors were relatively similar in size, as reflected in their total cell counts between 94,000 to 100,000 cells in the tumor bearing hemispheres (Fig 2D). To ensure data quality, we evaluated per-cell transcript density. The raw count distribution demonstrated robust capture efficiency, with a mean of 992.2 and a median of 823 transcripts per cell (Fig 2E). Subsequently, we used unsupervised clustering to group cells into 13 distinct clusters based on transcriptional similarity (Fig 3A, B). Clusters 0 and 1 contained the highest cell numbers (Fig S1A, B) and together formed the malignant tumor proper (Fig 3A). We then used a custom marker set to annotate these clusters by cell type (Fig 3C). Cluster 0 exhibited the highest expression of proliferation markers and was localized exclusively within the tumor region (Fig 3D); thus, we annotated this group as proliferative GBM cells (GBM_Prolif). Cluster 1 cells showed lower proliferation markers, but significant expression of genes involved in cytoskeletal rearrangement (Fig S1B); as their localization was also restricted to the tumor region, we annotated them as migratory GBM cells (GBM_Mig) (Fig 3E). We also identified two clusters (4 and 10) with strong macrophage/microglia gene signatures (Fig 3C), which differed significantly in their spatial localization. Cluster 4 cells were almost exclusive to the GBM region and were annotated as tumor-associated macrophages (TAMs) (Fig 3F). In contrast, Cluster 10 cells were found only in the non-tumor peripheral healthy tissue and were annotated as resident microglia (Fig 3G). Because initial unsupervised clustering did not provide sufficient resolution to isolate lymphoid populations, we employed a custom gating strategy to group CD4+ and CD8+ cells. The expression of T-cell-specific markers confirmed that this strategy yielded pure lymphoid populations (Fig 3C). Given the important role of iron in regulating immune dynamics in the TME, the current study focuses on the infiltration and interaction of the infiltrating myeloid and lymphoid populations with the two GBM cell clusters. The global localization of involved clusters (cell types) can be visualized in the UMAP graph (Fig 3H).

**Fig 2:**
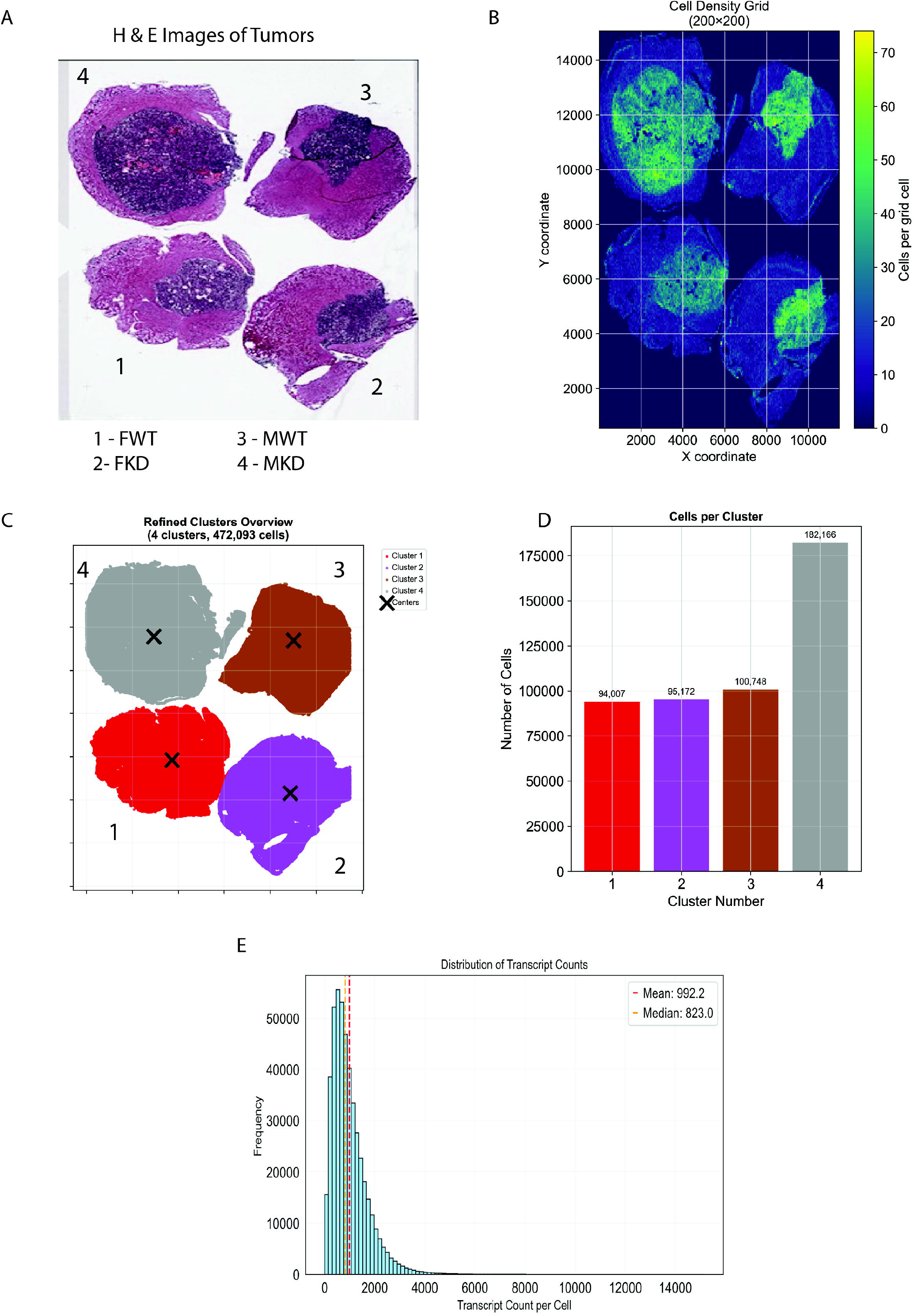
Characterization and segregation of the tumor bearing tissues according to their genotype. The four genotypes are Female WT (FWT; #1), Female FTH1 KD (FKD; #2), Male WT (MWT; #3), Male FTH1 KD (MKD; #4). (N = 1, Spatial transcriptomics) (A) H&E images of the four tumor bearing tissues used in Xenium analysis. (B) Cell density analysis showing a higher density of cells in the annotated tumor region of the brain tissues. (C) Clustering of the cells by their genotypes (D) Total cell counts per genotype showing that MKD (#4) has the highest number of cells. (E) Per-Cell transcript density showing the raw transcript counts as part of the Quality Control (QC) analysis.

**Fig 3:**
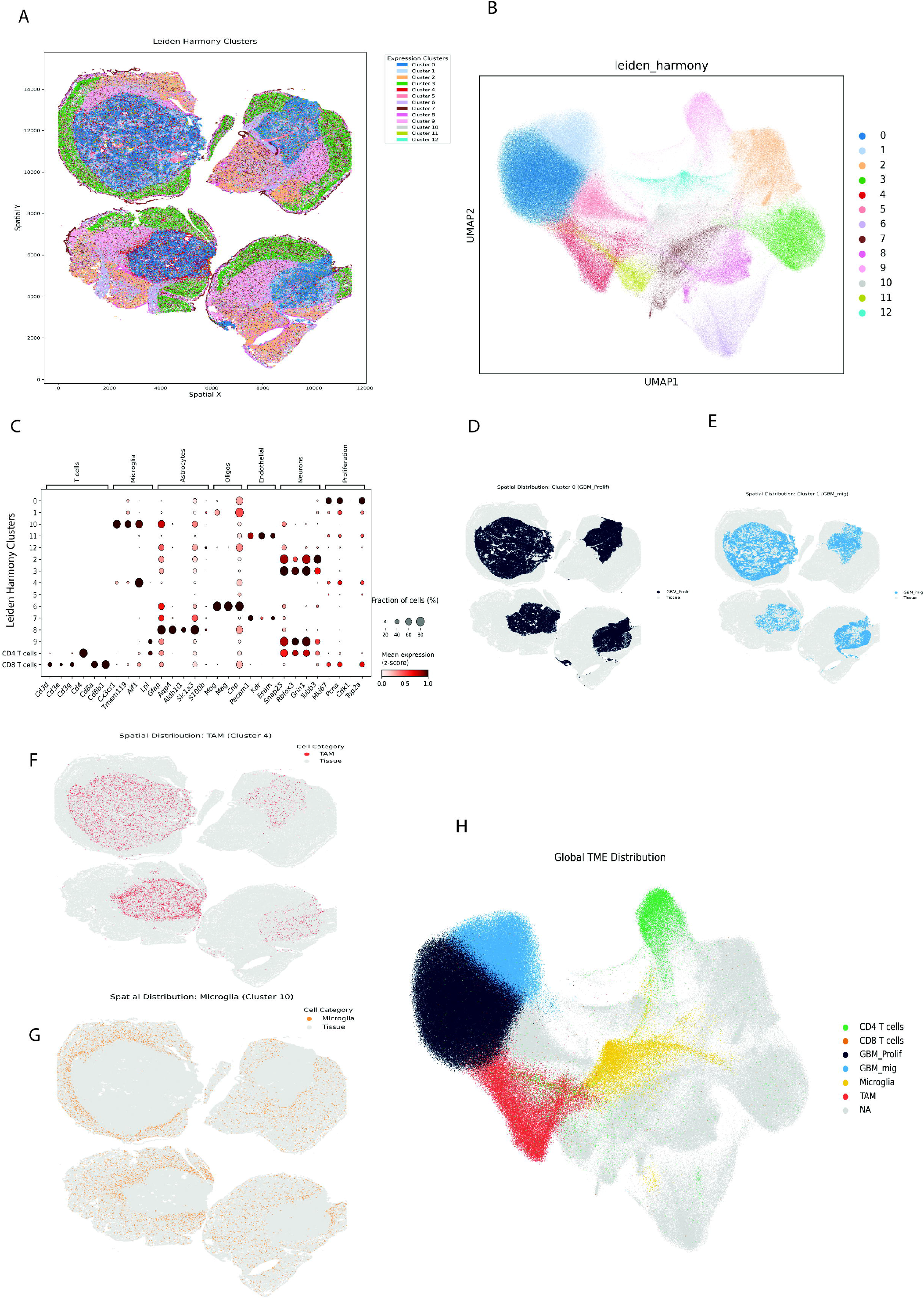
Clustering and characterization of cell types according to their transcriptional signatures. (N = 1, Spatial transcriptomics) (A) Demarcation of 13 leiden_harmony (expression) clusters across the four tumor bearing brains. (B) UMAP representation of the expression clusters. (C) Characterization and annotation of the clusters by custom set of cell type markers. (D, E) Localization of the GBM cell clusters (GBM_prolif – Cluster 0 and GBM _mig – Cluster 1) on the brain tissues. (F) Localization of the TAM cell cluster (Cluster 4) on the brain tissues, where they are exclusively present in the tumor region. (G) Localization of the microglia cell cluster (Cluster 10) on the brain tissues, where they are distributed throughout the non-tumor regions of the brain. (H) UMAP plot highlighting the GBM, T cell, TAM and microglia population.

### FTH1 KD reduces T cell infiltration into the TME in a chemokine driven manner

To investigate how extrinsic FTH1 knockdown (KD) affects T-cell recruitment, we utilized flow cytometry to quantify infiltrated T cells within the tumor-bearing hemisphere (FWT; N=8, FKD; N=9; MWT; N=11,MKD; N=10)^1^. In females, FTH1 KD significantly reduced CD4+ T-cell infiltration, whereas no significant difference was observed in males (Fig 4A). CD8+ T-cell levels followed a similar trend, with FTH1 KD reducing infiltration specifically in females (Fig 4B). Across both genotypes, male tumors exhibited lower baseline levels of CD4+ and CD8+ infiltration compared to female tumors (Fig 4A, B). We next interrogated T-cell infiltration within the four tumors used for our spatial transcriptomics analysis. To account for variations in tumor size, T-cell counts were normalized to the total number of GBM cells. Consistent with our flow cytometry results, FTH1 KD reduced both CD4+ and CD8+ infiltration into the TME in female mice. In males, we observed a reduction in CD4+ T cells, but no significant difference in CD8+ T-cell infiltration was detected (Fig 4C, D). Notably, both MWT and MKD tumors showed minimal CD8+ infiltration whether by flow cytometry or spatial transcriptomics. Finally, as T-cell infiltration is primarily driven by chemokine signaling [28], we quantified chemokine expression within the tumor regions of the four spatial tissues. In alignment with our infiltration data, female tumors showed a significant reduction in chemokines associated with CD4+ (Fig 4E) and CD8+ (Fig 4F) T-cell recruitment following FTH1 KD. Interestingly, two chemokines CCL17 and CCL20, showed increased expression in FTH1 KD female mice compared to WT. The discrete strong flip in expression of CD8+ cytokines also aligns with the severe diminution of CD8 T cells (Fig 4D). While some differences were noted in males, they were less pronounced. Overall, male tumors exhibited significantly lower baseline chemokine expression than female tumors, correlating with their lower T-cell infiltration. These data indicate that extrinsic FTH1 knockdown significantly impairs T-cell recruitment in female tumors possibly by decreasing TME chemokine expression. In contrast, male tumors are characterized by lower baseline chemokine levels and T-cell infiltration, which remain largely unaltered by FTH1 knockdown.

**Fig 4:**
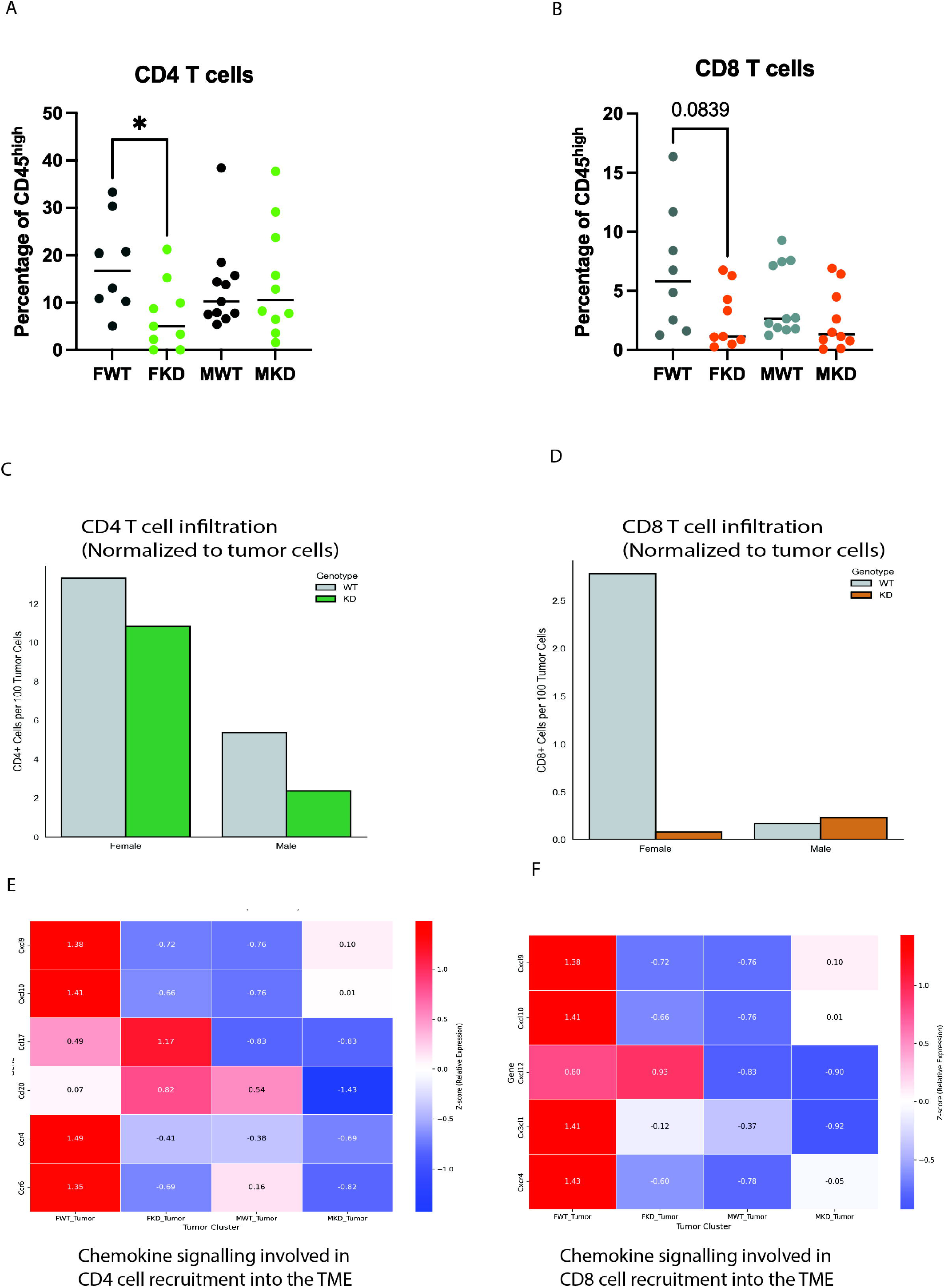
FTH1 KD reduces T cell infiltration into the TME in a chemokine driven manner. (A) In female but not male mice, FTH1 knockdown is accompanied by a decrease in CD4 T cell infiltration (FWT; N=8, FKD; N=9; MWT; N=11, MKD; N=10, One Way ANOVA). (B) In female but not male mice, FTH1 knockdown is accompanied by a decrease in CD8 T cell infiltration (FWT; N=8, FKD; N=9; MWT; N=11, MKD; N=10, One Way ANOVA). (C) After FTH1 knockdown, the abundance of CD4+ T cells in the tumor shows a 20% decrease in females and a 50% decrease in males. (N = 1, Spatial transcriptomics) (D) After FTH1 knockdown, the abundance of CD8+ T cells in the tumor shows pronounced female biased decrease. (N = 1, Spatial transcriptomics) (E) Heatmap depiction of a female biased shift in chemokine expression (z-score) involved in tumor-infiltrating CD4+ T cells after FTH1 knockdown. (N = 1, Spatial transcriptomics) (F) Heatmap depiction of a female biased change in chemokine expression (z-score) involved in tumor-infiltrating CD8+ T cells after FTH1 knockdown. (N = 1, Spatial transcriptomics)

### Ligand-Receptor (LR) interaction analysis shows sex biased remodeling of GBM-T cell interactions after extrinsic FTH1 knockdown

To further explore specific GBM-to-T cell signaling altered after extrinsic FTH1 knockdown, we utilized COMMOT analysis to compute the probability and signal flow of specific ligand receptor signaling pathways[25]. A physical distance threshold of 100 µm was applied to the spatial transcriptomic data, allowing the capture of both direct cell–cell LR interactions and indirect, secreted LR signaling (Fig 5A). We then separated GBM_Prolif–T cell and GBM_Mig–T cell interactions, treating GBM cells as senders and T cells as receivers. We observed distinct sex-specific gains and losses in signaling pathways related to T cell recruitment, adhesion, activation, and inflammation (Fig 5 B, C, D, E). In males, knockdown (MKD) tumors showed increased complement and BAFF signaling, consistent with a proinflammatory, innate immune–stimulating environment[29, 30]. In females, however, this phenotype was reversed, indicating suppression of inflammation and innate immune responses. Additionally, PTPRC was among the most significantly reduced signals in female knockdown (FKD) tissue. PTPRC (CD45) is an essential regulator of T cell activation and T cell receptor (TCR) signaling[31], and its loss is consistent with reduced T cell activation in FKD tumors. In males, the opposite pattern was observed, with PTPRC upregulated, supporting enhanced T cell activation. The most striking differences were seen in chemokine signaling. In males, FTH1 knockdown upregulated proinflammatory cytokine signaling, including IL6- and IL11-mediated pathways. In contrast, females showed a near-complete loss of T cell–recruiting CCL-, CCR5-, and CXCL-mediated signaling after FTH1 knockdown. Similar patterns were observed in both GBM_Prolif–T cell and GBM_Mig–T cell interactions, suggesting that these GBM subpopulations engage infiltrating T cells in comparable ways.

**Fig 5:**
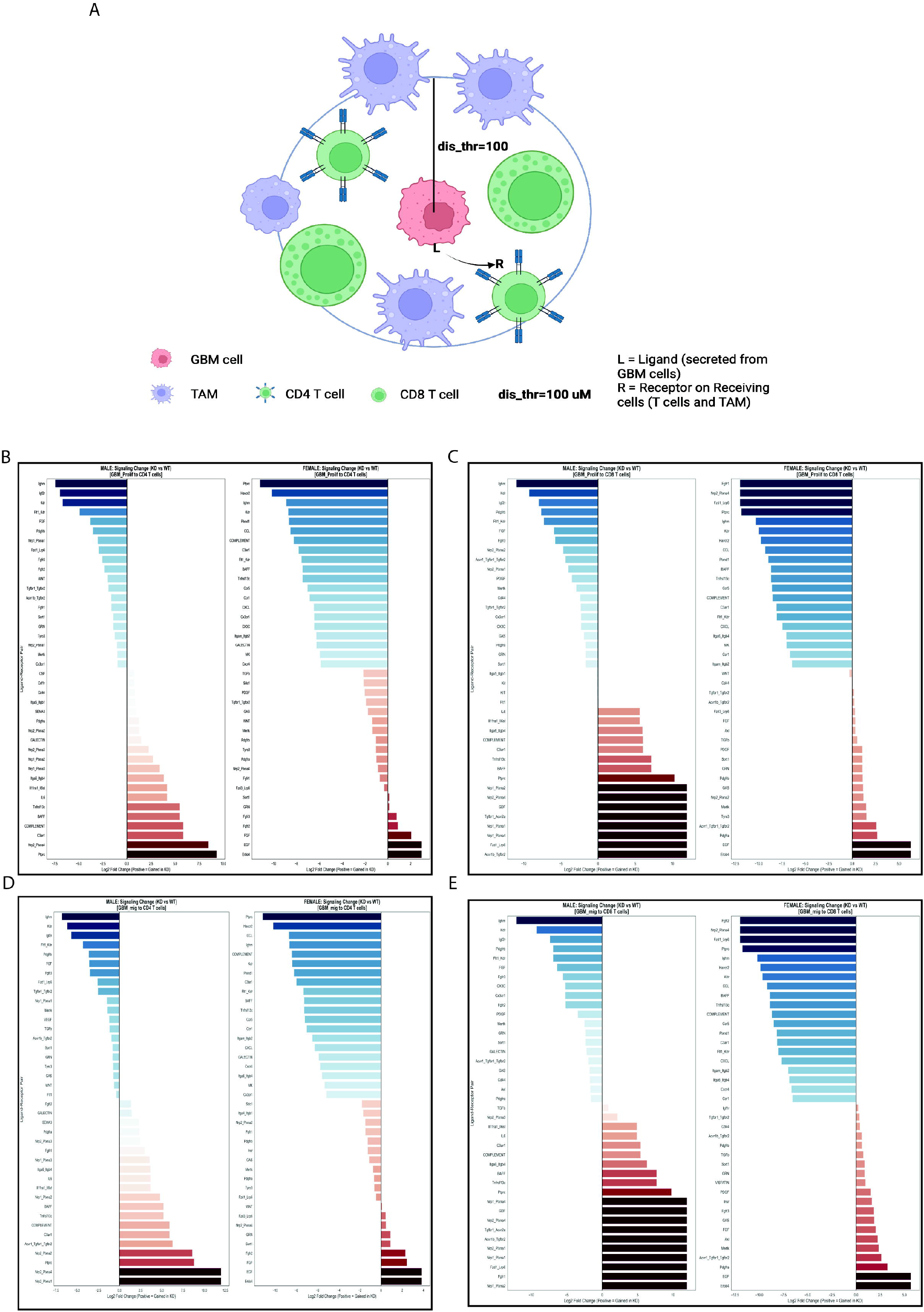
Ligand-Receptor interaction analysis between GBM cells (sender) and infiltrating T cells (receiver) (A) Graphical representation of COMMunication analysis by Optimal Transport (COMMOT) to characterize cell-cell communication between GBM cells and T cells where GBM cells act as senders and T cells as receivers, within a 100μm radius. (N = 1, Spatial transcriptomics) (B) The probability of ligand-receptor signals between GBM_Prolif and CD4 T cells. (C) The probability of ligand-receptor signals between GBM_Prolif and CD8 T cells. (D) The probability of ligand-receptor signals between GBM_Mig and CD4 T cells. (E) The probability of ligand-receptor signals between GBM_Mig and CD8 T cells.

### Extrinsic FTH1 knockdown alters TAM localization in GBM microenvironment

Iron availability in the TME has been shown to directly regulate myeloid cells infiltration. Flow cytometry analysis of the tumor bearing brain showed that FTH1KD tumors had elevated myeloid (CD11b+) cell infiltration in a female biased manner (Fig 6A). In GBM, majority of the infiltrating myeloid cells are represented by TAMs in human tissues and murine models. We utilized the spatial transcriptomic data to interrogate how these TAMs distribute in the tumor-bearing brains of the models employed. FTH1 KD increased the density of TAMs in the immediate periphery of the GBM cells (Fig 6B), in both males and females where females showed the highest difference. Next, to interrogate whether the two dichotomous cell states of GBM cells (proliferative and migratory) affect TAM distribution, we measured changes in TAM density relative to GBM_Prolif and GBM_Mig cells. FTH1 KD increased the local TAM density in females for both GBM_Prolif and GBM_Mig cells, however, in males only GBM_Mig cells exhibited higher TAM density (Fig 6 C, D). Finally, we interrogated the LR interaction present between GBM cells and TAMs by using cellchat[26]. Vcam1-(Itga4+Itgb1) signaling, which is involved in leukocyte recruitment and adhesion to the TME, showed higher probability in FTH1-KD mice and prominently towards GBM_mig cancer cells (Fig 6E, F). Similarly, the GBM_mig driven TAM interactions in MKD tumors show elevated signaling of genes involved in TAM adhesion and migration (Vcam1-Itgb, Sema4g-Plxnb2). These data explain the GBM_mig driven male specific upregulation of TAM density in the neighborhood (6E, F).

**Fig 6:**
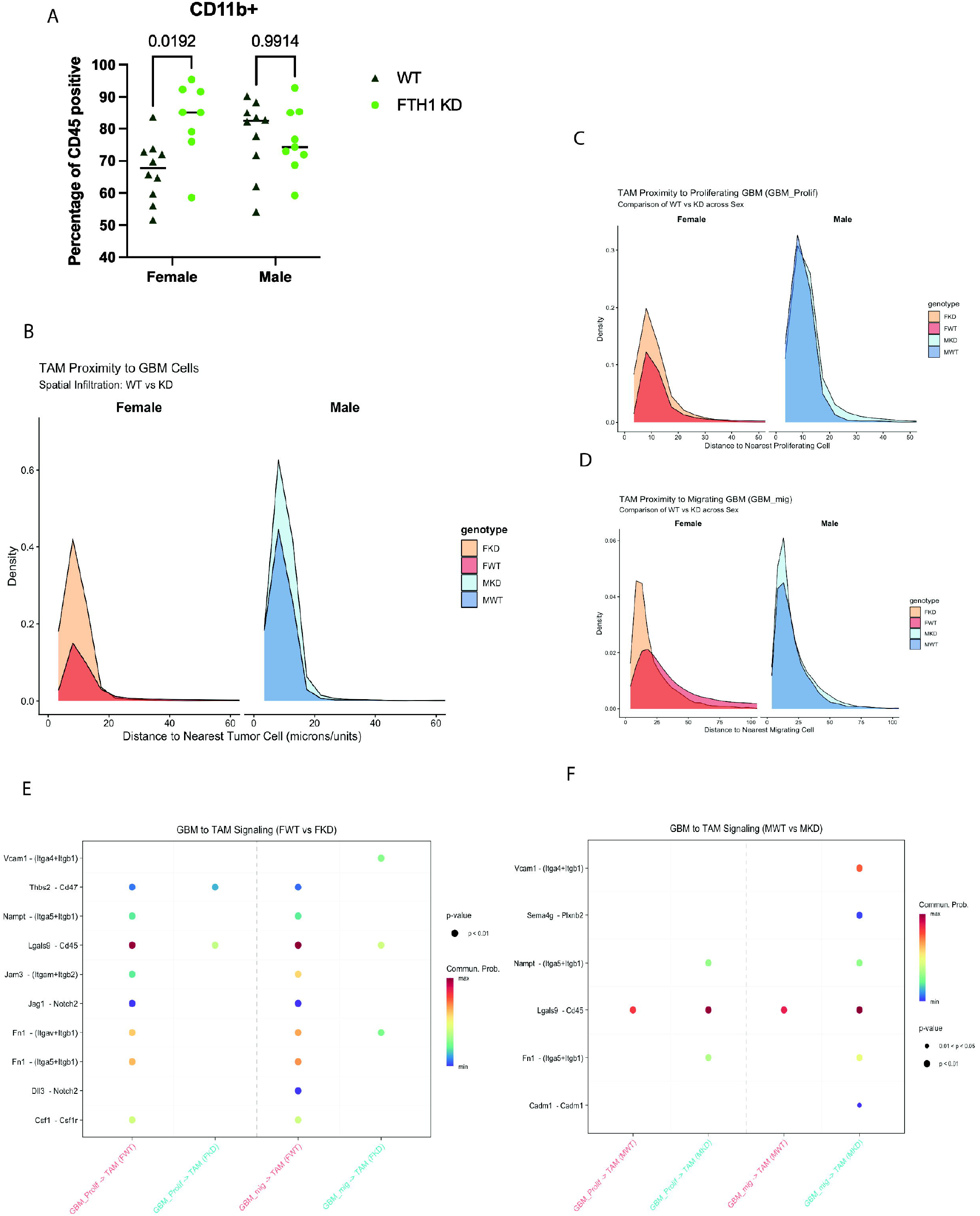
Extrinsic FTH1 knockdown alters TAM localization in GBM microenvironment. (A) In the FTH1 WT condition, tumors in males show higher infiltration by CD45+ CD11b+ cells than tumors in females. FTH1 KD has minimal impact in male hosts, but a pronounced increased infiltration in female mice. (One way ANOVA, MWT; N=10, MKD; N=9, FWT; N= 10, FKD; N=8). (B) Proximity analysis showing that FTH1 knockdown mice have higher density of TAMs near the GBM cells. (N = 1, Spatial transcriptomics) (C, D) Proximity analysis specific to GBM_prolif and GBM_Mig cells and TAM infiltration. (N = 1, Spatial transcriptomics) (E) CellChat analysis showing the gained and lost GBM-TAM signaling in females (FKD vs FWT). (N = 1, Spatial transcriptomics) (F) CellChat analysis showing the gained and lost GBM-TAM signaling in males (MKD vs MWT). (N = 1, Spatial transcriptomics)

## Discussion

Given iron’s essential role in tumor progression, its depletion has been considered a promising strategy to limit tumor growth. However, iron’s role in the tumor microenvironment (TME) is complex. On one hand, it is required for cancer cell proliferation; on the other, it is crucial for immune cell recruitment and function in mounting an antitumor response. Therefore, studying how TME dynamics change during iron deficiency can provide insights into therapeutically targeting iron metabolism to inhibit cancer growth. Our model provides a robust framework to study iron depletion in the GBM TME. Previous studies have shown that FTH1 can bind to GBM tissues[18]; however, our study is the first to definitively demonstrate that systemic FTH1 deficiency causes iron deficiency within the tumor. FTH1 KD mice exhibit decreased tumor iron levels, as measured by ICP-AES, along with tumor gene signatures indicative of cellular iron deficiency. IRP1/2 are cytoplasmic RNA-binding proteins that act as key post-transcriptional regulators of cellular iron metabolism. The tumors in FTH1 KD mice exhibit elevated IRP1/2 expressions, which is generally associated with cellular iron deficiency[32]. The tumors also show decreased expression of the iron exporter FPN. Depleted cellular FPN is a classic hallmark of iron deficient cells, as the cells actively try to inhibit FPN-mediated iron release to conserve cellular iron. HFE acts as a competitive inhibitor of TFR1 mediated iron uptake[33, 34]. Our results show decreased levels of HFE in the FTH1KD mice, which could be the result of a compensatory mechanism adapted by the GBM cells to elevate TFR1 mediated iron uptake. Previous studies have shown that cancer cells frequently decrease HFE expression to meet the high iron demand[7, 35]. Similarly, the decreased expression of cellular HJV in the tumor can be seen as a response to cellular iron deficiency. HJV is involved in the production of Hepcidin, a critical regulator of macrophage iron release[36]. Decreased HJV leads to decreased Hepcidin in the TME, which would promote macrophage iron release and tumor iron uptake.

Iron deficiency has been linked to changes in immune function and TME dynamics[19, 37, 38]. Iron is essential for T cell recruitment and activation[39], TAM polarization[40], migration and proliferation of tumor cells[41, 42]. Previous studies on this GBM model from our lab has shown that FTH1 KD affects T cell infiltration into the TME in a sex biased manner, where FKD tumors have lower T cell infiltration, corresponding with decreased survival[22]. We used spatial transcriptomics analysis to further interrogate the mechanism involved in this sex biased decrease of T cell infiltration. Here, we used unsupervised clustering of the cells by their transcriptional signature to distinguish 13 different cellular clusters. Of these clusters, cluster 0 and cluster 1 represent two transcriptional and functional states of GBM cells. Interestingly, we saw distinct proliferation and cytoskeletal rearrangement signature between the two tumor clusters, where cluster 0 was more proliferative in nature whereas Cluster 1 had higher expression of genes involved in cellular movement and cytoskeletal rearrangement. Hence, they were annotated as GBM_Prolif and GBM_Mig respectively for the subsequent analyses. Our results confirm the transcriptional heterogeneity in GBM cells which contribute to different cell states, as has been previously observed in GBM tissues[43-45].

Previous research has interrogated the role of abnormal cellular and tissue iron levels with impaired T cell infiltration and function [22, 46-48]. Both flow cytometry and spatial transcriptomics analysis showed a female biased decrease in T cell infiltration after FTH1 knockdown. Our data in WT mice also demonstrate the well-established sex-based differences in T cell infiltration, with male tumors exhibiting reduced CD8^+^T cell infiltration, consistent with previous studies. [49]. We have previously demonstrated that these FTH1 KD mice show no changes in circulating T-cell populations in peripheral blood, suggesting that the observed reduction in T-cell recruitment is specific to the GBM TME[22]. T cell infiltration into the tumor microenvironment (TME) is largely driven by chemokine signaling, with chemokines secreted by GBM cells actively recruiting CD4^+^and CD8^+^T cells. Our spatial transcriptomics analyses revealed that the reduced infiltration of CD4^+^and CD8^+^T cells was associated with a tumor-wide decrease in the expression of key chemokines (such as Cxcl9, Cxcl10, and Cxcl12) and chemokine receptors (including Ccr6, Ccr4, and Cxcr4) that regulate T cell recruitment. This is consistent with the depletion of T cells after FTH1 knockdown being chemokine driven. Additionally, Cxcl9 and Cxcl10 are known interferon-γ–inducible chemokines[50]. Iron acts as a key factor in IFN-gamma signaling, and iron deficiency can reduce IFN-gamma signaling in the TME[51]. Consequently, reduced interferon-γ activity may lead to decreased chemokine production, consistent with the tumor-wide chemokine profiles we observed. Interestingly, we observed a female specific upregulation of CCL17 and CCL20 in FTH1KD mice. CCL17 is a chemokine that is primarily secreted by M2 macrophages[52]. The iron deficient environment in the FTH1 KD female mice promotes M2 macrophage polarization, which leads to higher CCL17 secretion in the TME[53]. CCL20, on the other hand is mainly regulated by HIF proteins associated with hypoxia[54], and previous reports have established that iron chelation can lead to CCL20 secretion[55]. However, the spatial transcriptomic Z-scores represent a static endpoint measure of the tumor microenvironment at the time of tissue collection. Consequently, whether changes in chemokine abundance drive or reflect alterations in T-cell infiltration cannot be determined from the current data alone.

Chemokine signaling involves secreted cytokines (ligands) binding to their respective receptors in the target cells. In the current scenario, the GBM cells are the senders secreting chemokines into the TME. The chemokines then bind to cells expressing the chemokine receptor (TAMs, T cells). To further interrogate how these ligand-receptor (LR) signaling pathways are being affected, we used COMMOT (COMMunication analysis by Optimal Transport), a computational tool designed to infer cell-cell communication from spatial transcriptomics datasets [25]. We observed distinct sex biased changes in LR signaling after FTH1 knockdown. In males, FTH1 KD led to increased complement and its associated C3ar1 signaling between the GBM cells and the T cells. Complement signaling is normally associated with innate immune response and inflammation[56]. However, in T cells, complement signaling has been known to suppresses T-cell-mediated antitumor immunity[57]. Interestingly, there is an increase in BAFF signaling, involved in T cell activation and promoting T cell survival via activating the PI3K-Akt pathway in T cells[58]. The LR signaling data indicates that FTH1 KD induces a proinflammatory microenvironment in males as evidenced by increased BAFF, Complement and IL6 signaling. MKD tumors also lost FGF and associated FGFR signaling, which is involved in creating an immunosuppressive tumor microenvironment[59]. Combined, FTH1 KD in male mice drives a proinflammatory and less immunosuppressive tumor microenvironment. In females, FTH1 WT has high probability of chemokine mediated signaling between GBM cells and T cells, which is lost in FTH1 KD. These chemokine pathways include CCL, C3ar1, Ccr5, Ccr1, CXCL, Cx3cr1 and CX3C. The loss of these signaling in female FTH1 KD aligns with our GBM chemokine gene expression data and explain why the tumors in female FTH1KD mice have low T cell infiltration. Surprisingly, unlike males, FTH1 KD in the female host caused a decrease in proinflammatory (BAFF and Complement) signaling. Combined, FTH1 KD in females causes an anti-inflammatory, chemokine deficient tumor microenvironment that causes decreased T cell infiltration.

TAM recruitment and polarization can be affected by iron availability in the TME[60, 61]. We observed increased myeloid cell infiltration into the TME in FTH1 KD mice with a female bias. To further interrogate whether these infiltrating macrophages directly interact with the GBM cells, we measured the proximity of TAMs to GBM cells. We observed that FTH1 KD increased the density of TAMs in the immediate surrounding of GBM cells. Furthermore, the increased TAM density in the male FTH1 KD mice was mostly driven by the GBM_Mig cells. Finally, cellchat analysis[26] to characterize LR interaction between GBM cells and TAM provided insights into the heterogenous interaction of the two GBM subpopulations with the infiltrating TAMs. The GBM_Mig specific expression of VCAM (interacts with integrin) and Sema4g (interacts with Plexin-B2) in male FTH1 KD mice, could be the reason why only GBM_mig cells had higher TAM density in male FTH1 KD tumor with respect to male WT tumors. The above two pathways are essential of leukocyte infiltration and binding to the TME[62-64].

The GL261 syngeneic mouse model is considered highly immunogenic relative to other syngeneic GBM models and to the characteristically immunosuppressive microenvironment of human GBM, which may amplify the immune-mediated effects observed and bias the specific mechanisms by which FTH1 manipulation influences T-cell infiltration and survival [65]. Additionally, although 10X Xenium enables high-resolution spatial profiling, our analysis characterized gene expression across the bulk tumor region, assuming it as a transcriptionally homogeneous compartment. It is well established that GBM exhibits profound intratumoral spatial heterogeneity, with regions of hypoxia, active neovascularization, and invasive tumor cells at the tumor-brain interface each representing microenvironmentally distinct niches. Bulk averaging across these spatially disparate regions may therefore obscure focal chemokine gradients or immune exclusion zones that are mechanistically relevant to T-cell recruitment.

In summary, this study reveals that genetically induced tumor iron deficiency by extrinsic FTH1 knockdown alters the TME dynamics in GBM cells in a sex biased manner. In males, FTH1 knockdown induces a proinflammatory environment and increases the infiltration of TAMs into the neighboring environment of GBM cells. Whereas in females, FTH1 knockdown leads to an anti-inflammatory, cytokine deficient TME, which fails to recruit CD4 and CD8 T cells into the TME. Our results also explain the female biased reduction in overall survival from FTH1 knockdown we had seen in previous studies[22].

Female mice gain a survival advantage over male mice with GBM because of their robust anti-tumor immune response, as seen in the increased CD4 and CD8 infiltration rates in female WT tumors compared to male WT. However, FTH1 knockdown in females reduces the T cell infiltration to that of a similar level of male tumor, resulting in reduced anti-tumor immune response and significantly decreased survival. Our results further elucidate the importance of tumor iron levels in immune infiltration and anti-tumor immunity in GBM and establish the sexually dimorphic role of tumor extrinsic iron sources on GBM progression.

## Footnotes

1 All experimental groups were initiated with n = 12 mice. However, a few mice were removed from the study during the experimental period due to technical and humane endpoint criteria, resulting in unequal final group sizes.

## Notes

### Competing Interest Statement

The authors have declared no competing interest.

### Funding

- NCI-NIH: 5P01CA245705-05

