## Supplementary figures and images for "Iron-deficiency in the tumor microenvironment reprograms tumor-immune interactions in a sex biased manner"

### Sup Figure 1

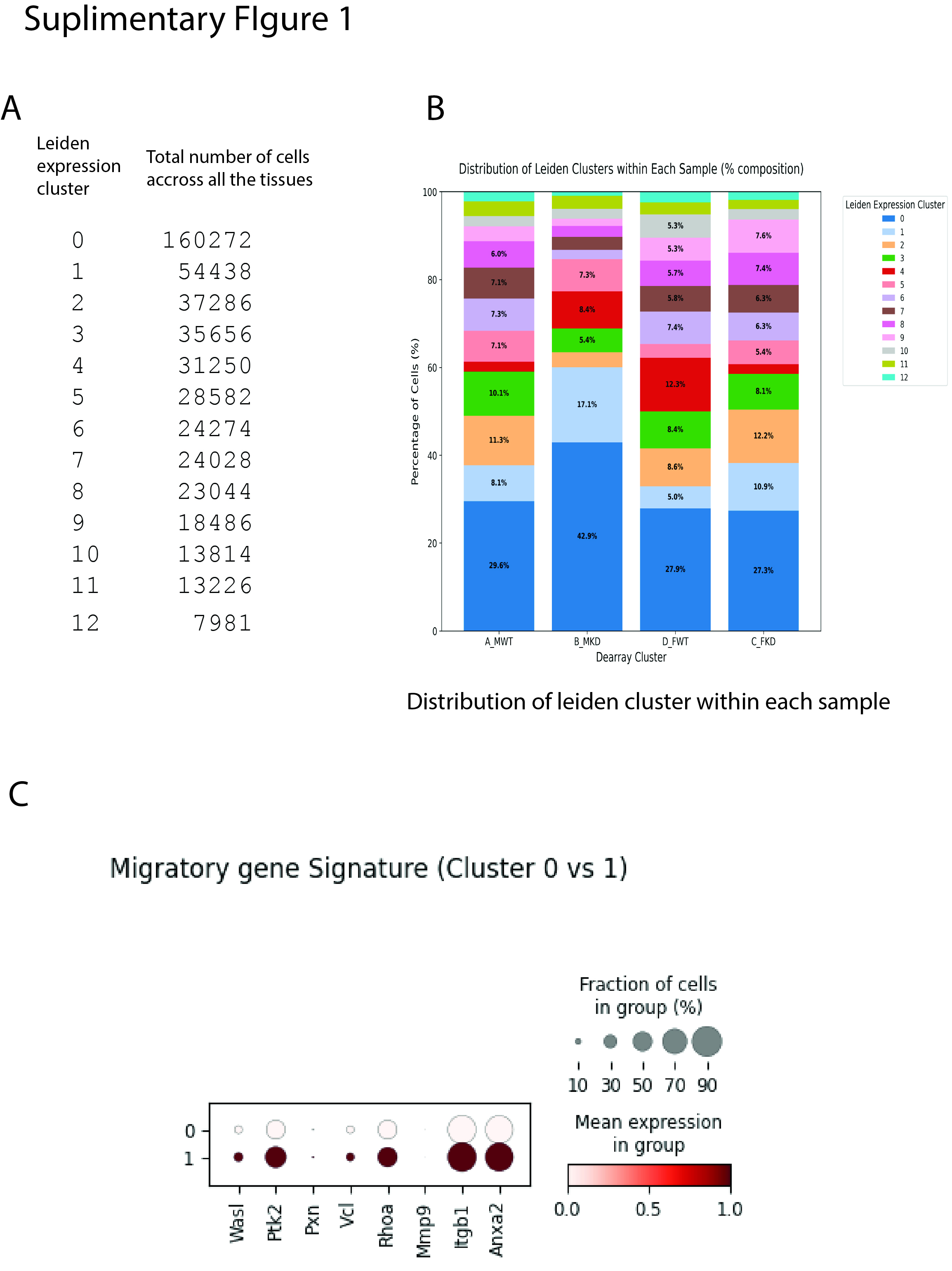
